# Blocking mitophagy does not improve fuel ethanol production in *Saccharomyces cerevisiae*

**DOI:** 10.1101/2021.06.29.450456

**Authors:** Kevy Pontes Eliodório, Gabriel Caetano de Gois e Cunha, Brianna A White, Demisha HM Patel, Fangyi Zhang, Ewald H Hettema, Thiago Olitta Basso, Andreas Karoly Gombert, Vijayendran Raghavendran

## Abstract

Ethanol fermentation is frequently performed under conditions of low nitrogen. In *Saccharomyces cerevisiae*, nitrogen limitation induces macroautophagy, including the selective removal of mitochondria, also called mitophagy. Shiroma and co-workers (2014) showed that blocking mitophagy by deletion of the mitophagy specific gene *ATG32* increased the fermentation performance during the brewing of Ginjo sake. In this study, we tested if a similar strategy could enhance alcoholic fermentation in the context of fuel ethanol production from sugarcane in Brazilian biorefineries. Conditions that mimic the industrial fermentation process indeed induce Atg32-dependent mitophagy in cells of *S. cerevisiae* PE-2, a strain frequently used in the industry. However, after blocking mitophagy, no differences in CO_2_ production, final ethanol titres or cell viability were observed after five rounds of ethanol fermentation, cell recycling and acid treatment, as commonly performed in sugarcane biorefineries. To test if *S. cerevisiae*’s strain background influences this outcome, cultivations were carried out in a synthetic medium with strains PE-2, Ethanol Red (industrial) and BY (laboratory), with and without a functional *ATG32* gene, under oxic and oxygen restricted conditions. Despite the clear differences in sugar consumption, cell viability and ethanol titres, among the three strains, we could not observe any improvement in fermentation performance related to the blocking of mitophagy. We conclude with caution that results obtained with Ginjo sake yeast is an exception and cannot be extrapolated to other yeast strains and that more research is needed to ascertain the role of autophagic processes during fermentation.

**Importance:** Bioethanol is the largest (per volume) ever biobased bulk chemical produced globally. The fermentation process is very well established, and industries regularly attain nearly 85% of maximum theoretical yields. However, because of the volume of fuel produced, even a small improvement will have huge economic benefits. To this end, besides already implemented process improvements, various free energy conservation strategies have been successfully exploited at least in laboratory strains to increase ethanol yields and decrease by-product formation. Cellular housekeeping processes have been an almost unexplored territory in strain improvement. Shiroma and co-workers previously reported that blocking mitophagy by deletion of the mitophagy receptor gene *ATG32* in *Saccharomyces cerevisiae* led to a 2.12% increase in final ethanol titres during Japanese sake fermentation. We found in two commercially used bioethanol strains (PE-2 and Ethanol Red) that *ATG32* deficiency does not lead to an improvement in cell viability or ethanol levels during fermentation with molasses or in a synthetic complete medium. More research is required to ascertain the role of autophagic processes during fermentation conditions.

## Introduction

High cellular viability and a good combination of product titre, yield and productivity are vital for the success of a bioprocess delivering bulk chemicals such as fuel ethanol. World bioethanol production is ca. 100 billion litres from corn, sugarcane, or wheat, using the industrial workhorse *Saccharomyces cerevisiae* (1, 2). The fermentation yield defined as grams of ethanol that can be obtained per gram of sugar is the most important parameter in this industrial process (3). In Brazil, decades of industrial practice and learning by doing have enabled the sugarcane biorefineries to achieve ∼85% of the theoretical maximum yield (0.511 kg of bioethanol per kg of hexoses) (4). Devising strategies to produce more fuel ethanol per hectare of land is challenging and requires nonconventional or breakthrough approaches. A small increase in yield using a robust yeast would represent enormous economic and environmental gains, due to the large volumes of bioethanol produced every year.

Bioethanol production can be described as a three-step process using large, non-aseptic tanks. Firstly, the sugarcane juice/molasses is fed into large tanks that contain the yeast slurry (prepared from the leftover of previous fermentations) and the mixture is allowed to ferment for ∼10 h; secondly, the fermented mixture is centrifuged to separate the yeast cells for use in the next cycle; and thirdly, the separated yeasts are washed with acid for 1-2 h to reduce the bacterial contamination, before they are utilised in a new fermentation cycle (5). Though the basic steps are common to all bioethanol plants, the yeast strains which catalyse the conversion of the sugars can be different in each biorefinery and accordingly several variants have been isolated (6). Basso and co-workers (6) investigated the population dynamics of yeasts in the industrial process to identify those strains that persist and dominate after several hundred rounds of cell recycling. They revealed that the baker’s yeast strains inoculated at the beginning of each sugarcane crushing season were naturally substituted by so-called wild or indigenous strains, some of which present many desired properties, such as absence of flocculation or foam formation. Very recently, it has been hypothesised that the biofuel producing yeasts may have been co-opted from a pool of yeasts preadapted to sugarcane juice fermentation in the cachaça industry, which in turn descend from wine yeast strains Europeans brought to South America, shedding new light on the origin and on the persistence and dominance of these wild yeasts (7).

Sugarcane juice/molasses do not have enough nitrogen for cell growth (8, 9). After a few division cycles, in response to nutrient depletion or stress, cells will consume part of their own contents in a process known as autophagy, to recycle nutrients, and in this process, some mitochondria are also degraded in a selective manner to maintain quality control (10–12). But minimal mitochondrial development is necessary for providing the cell with key metabolic intermediates and components (13). The role of mitochondria during anaerobic fermentation has not been given proper attention possibly because of glucose repression. In the presence of glucose, several respiratory enzymes and mitochondrial cytochromes remain low, vastly decreasing the occurrence of oxidative phosphorylation (14). Piggott and co-workers (15) used the genome wide gene deletion collection of the S288C strain to identify deletion strains with an advantage or a fitness defect over a 14-day period using synthetic grape juice. Strains with a defect in autophagy or in the ubiquitin-proteasome pathway had a reduced fitness indicating that genes that function in autophagy are required for optimal survival during fermentation even in a nitrogen replete environment (15).

Blocking mitochondrial degradation, known as mitophagy, improved cell viability and bioethanol yield, at least in the context of sake fermentation, which was achieved via disruption of the *ATG32* gene (16). When mitophagy is induced, the mitochondrial protein Atg32 binds to the adaptor protein Atg11, that links to the autophagosome biogenesis machinery. Sections of mitochondria are engulfed by growing autophagosomes that subsequently fuse with vacuoles. Inside the vacuole the mitochondria are then degraded (17, 18). To verify if an *ATG32* deficiency leads to better fermentation performance during Brazilian sugarcane fermentations for fuel ethanol production, the *S. cerevisiae* PE-2 strain, engineered to lack *ATG32*, was cultivated in a scaled-down mimicked industrial process, using molasses medium (5). Cells were assessed for their viability and bioethanol production over five rounds of cell recycling including sulphuric acid treatment. To compare our observations made using an industrial medium (molasses) and an industrial *S. cerevisiae* strain (PE-2), we also used the chemically defined, synthetic complete (SC) medium and carried out standard fermentation experiments over 10 days, with two additional *S. cerevisiae* backgrounds: the laboratory BY strain and the industrial yeast strain, Ethanol Red.

## Materials and Methods

### Microorganisms and maintenance

*Saccharomyces cerevisiae* PE-2 was obtained from NCYC; BY strains from EUROSCARF, and Ethanol Red™ was obtained from our in-house strain collection. Table 1 lists the genotypes and their origin. Strains were stored as YPD + glycerol (15% v/v) stocks (–80 °C). Strains were routinely grown on YPD (Yeast extract 10 g·L^-1^, Peptone 20 g·L^-1^ and Dextrose 20 g·L^-1^). When solid medium was required agar (20 g·L^-1^) was added. Routine growth was performed at 30 °C, 200 rpm.

**Table 1:**
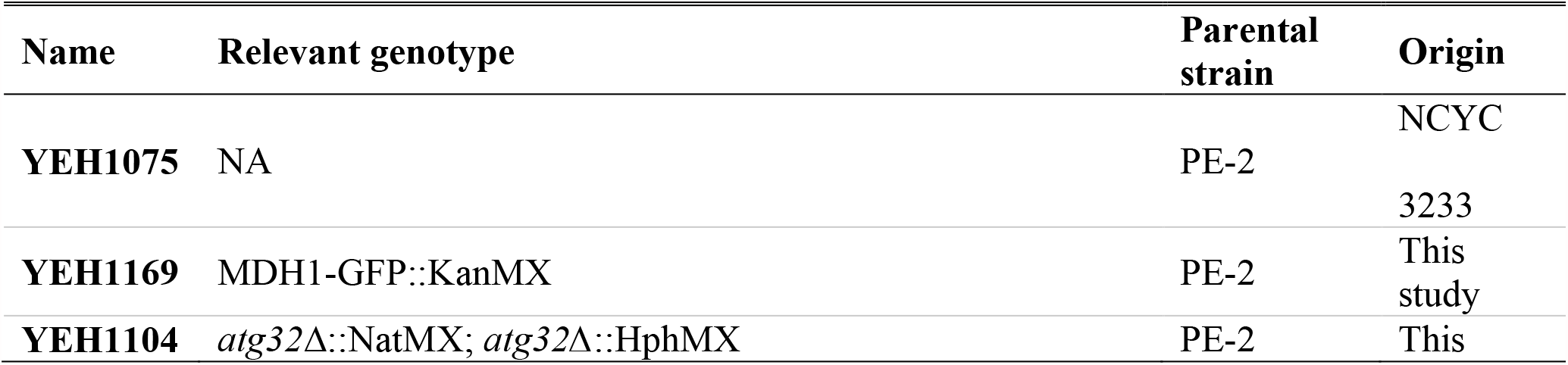
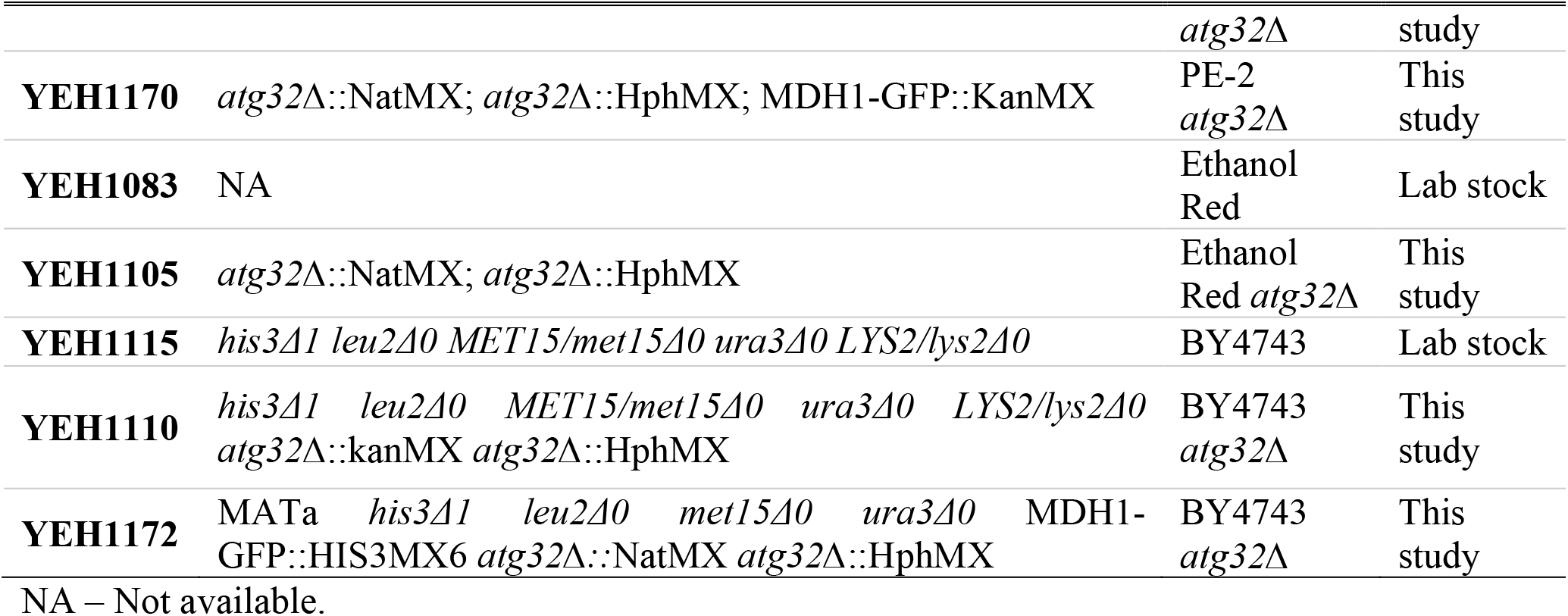
List of strains used in this study.

### Molecular biology techniques

Amplification of DNA cassettes were carried out using MyTaq DNA polymerase (Bioline Reagents Ltd, UK) in a thermocycler with desalted oligonucleotide primers (Sigma-Aldrich, UK) according to the manufacturer’s protocol. *S. cerevisiae* was transformed with the lithium-acetate method (19). Primers used in this study are listed in Table S1 (Supplementary material). Single-cell lines of transformants were obtained by re-streaking on solid selective medium.

### Strain construction

Two copies of the *ATG32* gene were mutated in Ethanol Red and PE-2 by replacing them with selectable markers using a PCR based gene deletion method. The first copy of the *ATG32* open reading frame was replaced by the *Klebsiella pneumoniae* hygromycin B phosphotransferase gene cassette (HphMX) that confers resistance to hygromycin B (20). This cassette, present on pAG32, was amplified using primers VIP3915 and VIP3916 (Table S1). Primers used for amplification contained 50 bp flanks identical to the *ATG32* ORF upstream and downstream region to enable gene deletion via homologous recombination. Transformed cells were plated onto YPD agar plates containing hygromycin B at a concentration of 300 µg·mL^-1^. The NatMX cassette on plasmid p4339 (21) was amplified using the same primers to replace the second *ATG32* ORF. NatMX confers resistance to nourseothricin (ClonNat, Werner Bioagents). Transformants were selected on YPD agar plates containing 100 µg·mL^-1^ ClonNat. Transformants were restreaked on selective medium and total DNA was isolated from overnight YPD cultures using glassbeads/phenol-chloroform extraction (22). Gene deletions were confirmed by PCR on total DNA. The plasmids used in this study are mentioned in Table S2 (Supplementary material).

Deletion of *ATG32* in the BY4743 strain was carried out in the haploid backgrounds (BY4741 and BY4742) using NatMX and HphMX as selection markers. Diploids were created by mating the strains and selecting on YPD plates containing both ClonNat and hygromycin B.

To visualise mitochondria, Mdh1 was C-terminally tagged with GFP using a one-step PCR-mediated protocol. The PCR fragment was generated using the primers VIP4045 and VIP4046 using pFA6a-GFP(S65T)-kanMX and pFA6a-GFP(S65T)-HIS3MX6 as template for PE-2 strains and BY4743 strains, respectively (23).

### Inoculum preparation

A loopful of cells from a single colony on solid YPD-agar was inoculated into YPD (10 mL) and grown for 17 h in an orbital shaker at 200 rpm at 30 °C. Cells were counted in a Neubauer chamber using a phase contrast microscope at 400× magnification. The required volume of culture liquid – corresponding to an initial optical density (600 nm) of 1.2 (Jenway 6300 spectrophotometer) at the start of the propagation – was centrifuged, pellet resuspended in sterile YPD or sterile synthetic complete medium (SC) (1 mL) and used as the inoculum for batch fermentation.

### Fermentation experiments

The medium and the experimental set up used in this study are summarised in table 2.

**Table 2:** Experimental conditions used in this study.

| Cultivation method | Setup | Medium + culture conditions |
| --- | --- | --- |
| 1 | 25/50 mL Erlenmeyer flask stoppered with Suba-Seal® | YPD, oxygen restricted, shaken, 30 °C, 175 rpm |
| 2 | 50 mL plastic centrifuge tube, loosely capped | Molasses, oxygen restricted, static, 32 °C |
| 3 | 50 mL plastic centrifuge tube, loosely capped | Synthetic complete medium, oxygen restricted, static, 30 °C |
| 4 | 8 mL Corning glass tubes loosely plastic capped/ cotton stoppered 250 mL Erlenmeyer flask | Synthetic medium, oxygen restricted, 30 °C, static/Synthetic complete medium, 30 °C, 150 rpm |

### Fermentation with rich complex laboratory medium

Fermentation was carried out as described by Argueso and co-workers (24) on modified YPD (yeast extract 5 g·L^-1^, peptone 10 g·L^-1^ and 100 g·L^-1^ glucose) in 25 mL Erlenmeyer flasks (cultivation method 1) containing 15 mL of medium and stoppered with a Suba-Seal® (Sigma Aldrich Inc.) septum to maintain oxygen restricted conditions. Cells were grown for two cycles in the same medium to provide an adaptation round before using them as inoculum for the fermentation experiments at a starting cell concentration of 10 mg_wet cells_·mL^-1^. Septa were pierced with two needles, one for sampling using a syringe, and another one for the gas exit. Flasks were shaken at 175 rpm in an orbital shaker at 30 °C to keep the cells suspended. *∼* 0.5 mL samples were taken using a needle and a syringe every three hours, centrifuged (12,000*g*, 4 °C), supernatant stored at –20 °C, and wet pellet weighed to monitor the increase in cell mass over time.

### Fermentation mimicking the Brazilian sugarcane biorefinery

Fermentation mimicking the Brazilian fermentation process was carried out as described previously (5) using molasses from São Manuel mill (São Paulo, Brazil) as a substrate (cultivation method 2). Briefly, 100 g·L^-1^ of molasses-grown wet cells were used to initiate the fermentation using molasses containing total reducing sugars (TRS) of 140 g·L^-1^. Approximately 4 g of wet cells were added to 50 mL centrifuge tubes and 27.75 mL sugarcane molasses was added in three equivalent parts at 0, 2, and 4 h of fermentation, mimicking the fed-batch process. The mass of CO_2_ lost was used as a proxy for the progress of the fermentation. The temperature was controlled at 32º C in a static incubator, conditions typically encountered during industrial fuel ethanol production. After 10 h, tubes were taken out of the incubator and left on the bench at room temperature. After 20 h, tubes were mixed well, and 1 mL sample was taken for cell viability and metabolites analysis. The tubes were then centrifuged, and the pelleted cells were resuspended in 6 mL of water, and 2 mL of the wine (supernatant) from the previous cycle, and subjected to acid treatment (H_2_SO_4_, 2 M) for 1 h to a final pH of ∼ 2.5 under static conditions before feeding was commenced with fresh molasses (5).

### Fermentation with mineral synthetic medium

Mineral synthetic medium used in fermentation tests consisted of yeast nitrogen base (Difco™) without amino acids (1.7 g·L^-1^), supplemented with ammonium sulphate (5.0 g·L^-1^), complete supplement mixture Drop-out: Complete (Formedium, UK) (790 mg·L^-1^), and anhydrous glucose (150 g·L^-1^) in deionised water. Glucose was autoclaved separately. Batch sterilisation was carried out using the benchtop autoclave using factory settings. Before the start of the fermentation, 40 mL of SC medium were aliquoted in 50 mL Falcon tubes, (Sarstedt), the tubes were sparged with N_2_ gas (BOC, UK, Oxygen Free Nitrogen, grade 5.0) for 2 min using sterile needles (10 cm length) and inoculated with the cell suspension for a start OD_600_ of 1.2. The tube caps were kept slightly loose to allow gas exit during the fermentation and maintained under static conditions at 30 °C (cultivation method 3). Tubes were weighed at time zero and then weighed every 24 h for 10 days to monitor mass loss which is used as a proxy for the progress of the fermentation.

### Fermentation kinetics in synthetic medium

In the experiment described in the above section, only end point sampling was made. To study the kinetics of cell viability during the fermentation, another set of experiments was conducted exactly as described above, but in 8 mL screw capped glass tubes (cultivation method 4). Cells were inoculated to ten 8 mL screw capped glass tubes containing 6 mL of SC medium (previously sparged with N_2_) and inoculated at the same OD as described above. One tube was taken out each day and samples were taken for cell count and metabolites analysis. Aerobic cultivations using SC medium were done in cotton stoppered 250 mL Erlenmeyer flasks (cultivation method 4) containing 50 mL medium and shaken at 150 rpm in an orbital shaker. Samples (1 mL) were taken every 24 h for cell count and metabolites determination.

### Mitophagy assays

Under respiratory conditions, growth on a non-fermentable carbon source leads to the proliferation of mitochondria. Thus, cells were grown overnight on YPglycerol (Yeast extract 10 g·L^-1^, Peptone 20 g·L^-1^ and glycerol 30 g·L^-1^) in long glass test tubes with plastic caps and shifted to nitrogen starvation medium (Dextrose 20 g·L^-1^, yeast nitrogen base (Difco™) without amino acids and ammonium sulphate. During starvation, a substantial fraction of mitochondria are sequestered as cargoes and transported to the vacuole. Fluorescent marker proteins used to monitor mitochondria can temporarily withstand the acidic conditions in the vacuole and can be detected by immunoblotting as semi-quantitative evidence for mitophagy (25).

For mitophagy assays on molasses (organic blackstrap molasses, Holland & Barrett, Meridian, UK), cells were grown overnight on YPD and shifted to molasses (140 g_total-sugars_·L^-1^) to an OD_600_ = 1.2 and incubated up to 44 h. From this culture, 1.3 g of wet cell pellet was transferred to 13 mL fresh molasses in a 15 mL falcon tube and incubated under oxygen restricted conditions for 8 h and 24 h at 30 °C. Cells were harvested and washed with water. Some particulate debris was observed in the molasses grown cells due to the impurities present in the molasses.

### Fluorescence imaging

Live cells were analysed with an Axiovert 200M microscope (Carl Zeiss, Inc. Oberkochen, Germany) equipped with Exfo X-cite 120 excitation light source (Excelitas technologies, Waltham MA, USA), band-pass filters (Carl Zeiss, Inc., and Chroma, Bellow Falls VT,USA), and a PlanFluar 100×/1.45 NA objective lens (Carl Zeiss, Inc.) and a digital camera (Orca ER; Hamamatsu, Hamamatsu City, Japan). Image acquisition was performed using Volocity software (PerkinElmer, Beaconsfield, UK). Fluorescence images were routinely collected as 0.5 µm Z-Stacks and merged into one plane using Openlab (PerkinElmer, Beaconsfield, UK) and processed further in Photoshop (Adobe) where only the level adjustment was used. Bright-field images were collected in a single plane and modified, so that only the circumference of the cell was visible and pasted into the blue channel of Photoshop.

### Western blot analysis

For preparation of cell extracts by alkaline lysis, 10 OD_600_ units of cells were harvested, and pellets resuspended in 0.2 M NaOH and 0.2% (v/v) β‐mercaptoethanol and left on ice for 10 min. Soluble protein was precipitated by addition of trichloroacetic acid (5% final concentration (w/v) and left on ice for a further 10 min. Following centrifugation (13,000*g*, 5 min, 4 °C), the pellet was resuspended in 10 μL of 1 M Tris–base and incubated at 95 °C in 90 μL 1× SDS–PAGE sample loading buffer for 10 min. Samples (0.5 OD_600_ equivalent) were resolved by 10% SDS–PAGE and transferred (BioRad Mini-Protean gel and transfer system) to 0.45 µm Nitrocellulose filters. Blots were blocked in 2% (w/v) fat‐free Marvel™ milk in TBS‐Tween 20 (50 mM Tris–HCl (pH 7.5), 150 mM NaCl, 0.1% (v/v) Tween 20). Proteins were detected using monoclonal anti‐GFP (mouse; 1:3000; Roche 11814460001). Secondary antibody was HRP‐linked anti‐mouse polyclonal (goat; 1:10000; Bio‐Rad, STAR207P). Detection was achieved using enhanced chemiluminescence (Biological Industries, Kibbutz Beit Haemek, Israel) and chemiluminescence imaging using a Gbox from Syngene (Synoptics Ltd, Cambridge, UK).

### Cell viability determination

For experiments in SC medium, cellular viability was measured by plating out 100 µL of diluted samples on YPD (1:50, 1:1000, 1:5000 dilutions) and incubating them at 30 °C for 48 h. Colonies were manually counted to calculate the viable colony forming units (CFU) per mL. For experiments with molasses, counting of live/dead cells was performed as described elsewhere (26).

### Flow cytometry analysis

The forward scatter channel (FSC) and side scatter channel (SSC) measurements at the end of each fermentation cycle of mimicked industrial fermentations were determined by flow cytometry. Flow cytometry analysis was performed by taking a 100 µL sample from each tube after fermentation had ceased (20 h). Samples were diluted 1000-fold in phosphate buffers saline (pH 7.4), resulting in a final cell concentration of ∼10^6^ cells·mL^-1^. The intensity of the FSC and SSC readings were obtained using CytoFLEX flow cytometer (Beckman-Coulter, USA), followed by post-acquisition performed in CytExpert analysis software (Beckman-Coulter, USA). Cell counting was made with 30 µL (∼10^5^ cells·mL^-1^.) of each yeast suspension.

### Extracellular metabolites determination

Glucose was analysed using the Megazyme GOPOD kit (K-GLUC 10/15, Megazyme Inc., Ireland); ethanol was analysed using the Megazyme LQR kit (K-ETOH LQR, Megazyme Inc., Ireland) using the manual assay procedure for large volumes (scaled down to half the volume suggested in the manual) in a semi micro-cuvette. Substrate and metabolite concentrations from molasses fermentation were determined by HPLC (Shimadzu Prominence, Kyoto, Japan) using a refractive index detector. Samples were centrifuged and filtered through 0.22 µm pore size filter prior to injection. Sucrose, fructose, and glucose in samples from cultivations with molasses were separated with an Aminex® HPX-87C ion-exchange column (Bio-Rad) at 85 °C, with ultrapure water as the mobile phase, at a flow rate of 0.6 L·min^−1^. Ethanol and glycerol were analysed with an HPX-87H ion-exchange column (BioRad) at 60 ºC with sulphuric acid (5 mM) at a flow rate of 0.6 L·min^-1^.

### Calculation of the maximum specific growth rate

The maximum specific growth rate was obtained by plotting the natural logarithm of the culture absorbance against time. The slope of the linear regression line represents the μ_max_.

### Statistical calculations

Two sample *t*-test for assuming unequal variances (95% confidence) was carried out in Microsoft Excel 365 using the data Analysis Toolpak add-in (Microsoft Inc.).

## Results

### Atg32 deficiency leads to the loss of mitophagy in the industrial strain *S. cerevisiae* PE-2

Nitrogen starvation in laboratory yeast strains induces the selective autophagy of mitochondria. During this process mitochondria are delivered to and degraded in vacuoles (10). To monitor mitophagy we made use of the observation that GFP fusion proteins are degraded upon entry into the vacuole but, since GFP is relatively resistant to vacuolar degradation, GFP temporarily accumulates after delivery to the vacuole and its accumulation can be used as a qualitative measure of autophagy (27). We tagged the mitochondrial malate dehydrogenase (Mdh1) with GFP in the genome of *S. cerevisiae* PE-2 (strain YEH1169, Table 1). These cells were grown on a nonfermentable carbon source (glycerol) before transfer to low nitrogen, glucose medium. Upon shift to nitrogen starvation, we observed the expected faint staining of a large subcellular structure, typical of the vacuole and a reduction in the extensiveness of the mitochondrial network (Fig. 1A). These changes are dependent on Atg32 and are in agreement with previous observations for S288c-derived strains (17, 18). In addition, we also monitored Mdh1-GFP entry into vacuoles by western blot analysis using antibodies against GFP. Mitochondrial GFP fusion proteins that enter the yeast vacuole are cleaved and a relatively protease resistant GFP fragment accumulates temporarily that can be detected with a GFP antiserum (10). Upon nitrogen starvation, the typical GFP fragment indeed becomes apparent in PE-2 cells expressing Mdh1-GFP but not in cells deficient of Atg32 (Fig. 1B). We conclude that Atg32 is required for mitophagy under nitrogen starvation in the industrial strain PE-2.

**Figure 1:**
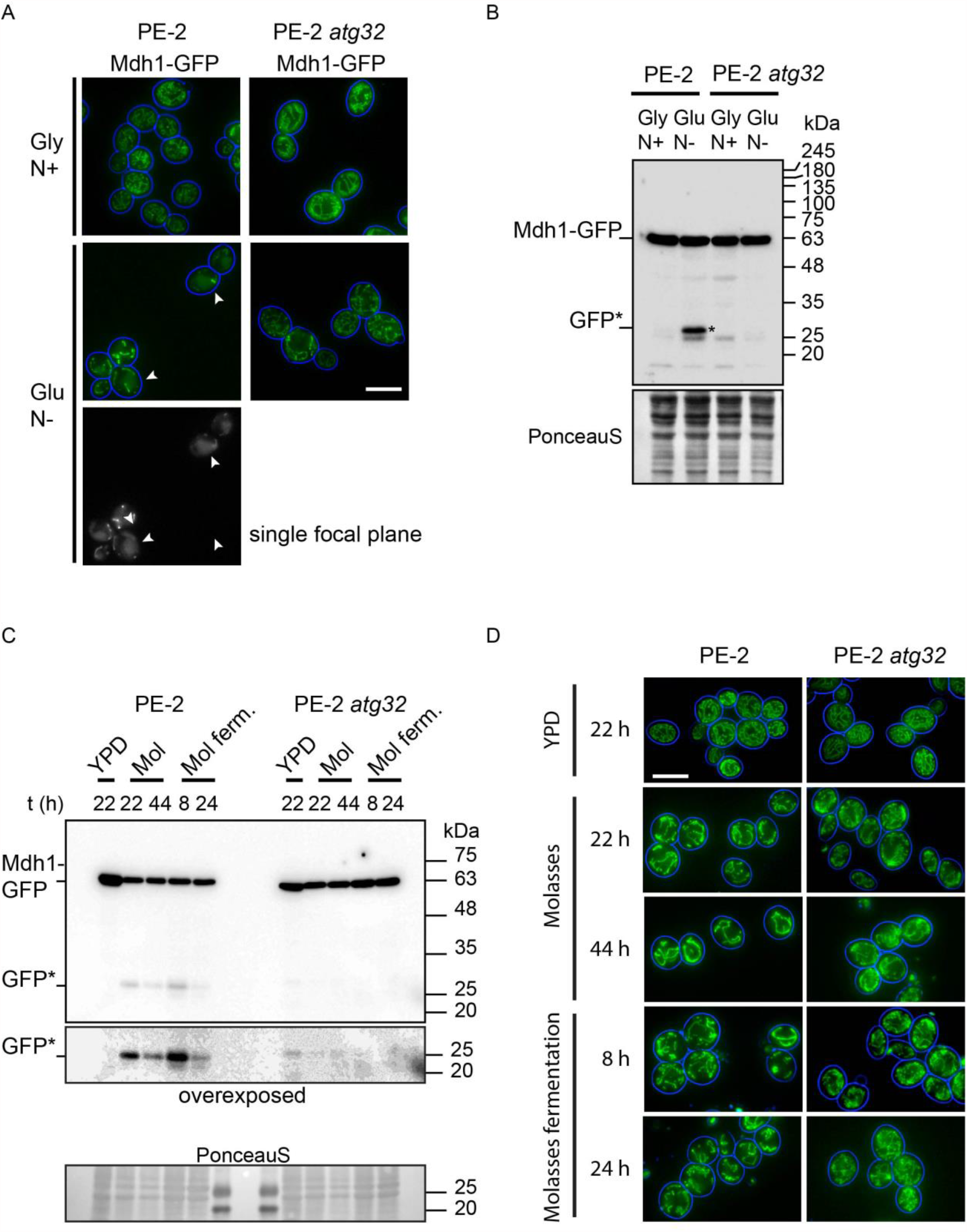
Mitophagy is blocked in *S. cerevisiae* PE-2 cells upon *ATG32* deletion. Mitophagy was assayed in *S. cerevisiae* PE-2 expressing Mdh1-GFP and isogenic *atg32* cells grown overnight on nitrogen-rich glycerol medium (Gly N+) and shifted to glucose, nitrogen starvation medium (Glu N-) and incubated for 6 h at 30 °C (A, B), or overnight on YPD and shifted to molasses and incubated up to 44 h. From this culture, cells were transferred to fresh molasses and incubated under oxygen restricted conditions (molasses fermentation, mol. ferm.) for 8 h and 24 h at 30 °C (C, D). Cells were imaged using epifluorescence microscopy and Z stacks were merged into a single green image. Cell circumference is stained blue (A, D). Note that upon nitrogen starvation (A), mitochondrial structures are less abundant in PE-2 cells and instead a large diffusely stained organelle becomes apparent (white arrow heads, vacuoles). A single focal plane near the centre of the cell is shown to clearly illustrate the vacuolar staining. Although mitochondrial structures are reduced in PE-2 cells in molasses compared to PE-2 *atg32* cells, no vacuolar labelling is observed. Scale bar, 5 µm. (B, C) Western blot analysis of PE-2 and PE-2 *atg32* cell samples using anti-GFP. GFP* indicates the relatively protease‐resistant GFP-degradation product indicative of vacuolar breakdown. An overexposed section of the blot around GFP* is included (C). A Ponceau S staining of the filter was used as loading and blotting control (B, C)

Subsequently, we tested whether PE-2 cells growing in molasses, which is known to be low in nitrogen, induce mitophagy. Both in the presence of oxygen and during fermentation, cells clearly display Atg32-dependent mitophagy (Fig.1C), albeit at a low level as no clear vacuole labelling of Mdh1-GFP could be observed, but the mitochondrial network remained much more extensive in PE-2 *atg32Δ* cells compared to PE-2 cells (Fig. 1D).

### Deletion of the mitophagy receptor *ATG32* did not affect fermentation in rich medium

To investigate if deleting the gene for a key cellular maintenance process affected growth under nutrient abundant conditions, PE-2, Ethanol Red, and BY wildtype (WT) and *atg32Δ S. cerevisiae* were grown in standard YPD medium (see inoculum preparation) but using a higher glucose concentration (100 g·L^-1^) under oxygen restricted conditions (cultivation method 1). No difference was observed in the maximum specific growth rate or in the final ethanol titres (Figure 2a and Figure 2b) between the WT and the *atg32Δ* mutants during a batch fermentation process. Ethanol Red exhibited the highest µ_max_ (based on OD_600_) (WT: 0.44 ± 0.01 h^-1^; *atg32Δ* 0.45 ± 0.00 h^-1^), followed by PE-2 (0.41 ± 0.01 h^-1^ for WT and *Δatg32*), and BY (0.37 ± 0.00 h^-1^ for WT and *atg32Δ*). We further investigated the kinetic performance of the *S. cerevisiae* PE-2 strain and its corresponding *atg32Δ* mutant in a similar medium than the one described above, but with less nitrogen (0.5× YP, cultivation method 1), also with 100 g/L initial glucose, after acclimatising the cells for two days aerobically (24). The results show that PE-2 WT and *atg*32*Δ* behaved identically in terms of biomass formation (Fig. 3a), sugar consumption and ethanol formation (Fig. 3b), reaching a final ethanol concentration of 50 g·L^-1^ in 12 h (Fig. 3b).

**Figure 2:**
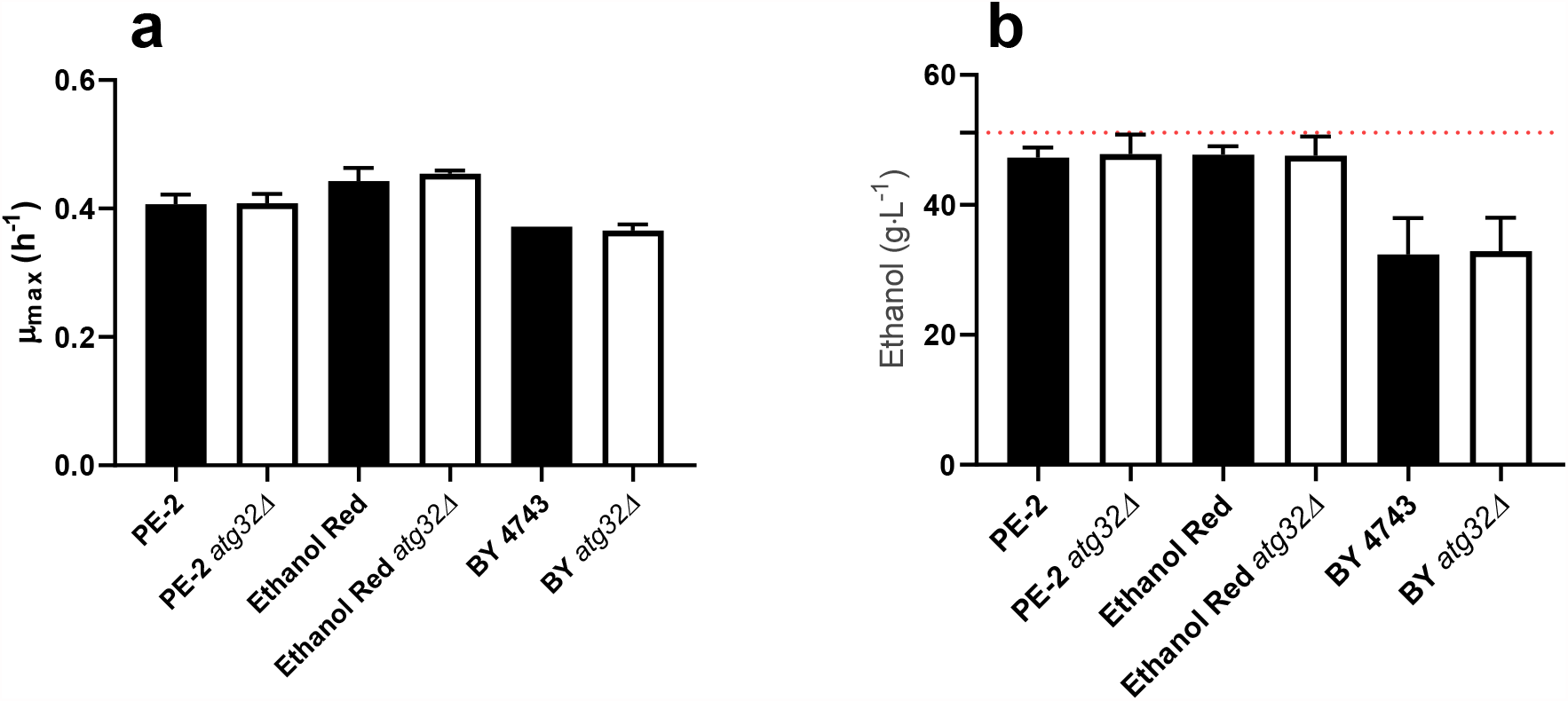
**a**) The maximum specific growth rate (µ_max_) and **b**) final ethanol titres during oxygen restricted growth in YPD using 100 g·L^-1^ initial glucose. The dashed line in 1b represents the maximum ethanol concentration that could be theoretically achieved. Filled columns represents the WT and open columns represents the *atg32 S. cerevisiae*. Data represent the mean and deviation from the mean (n = 2).

**Figure 3:**
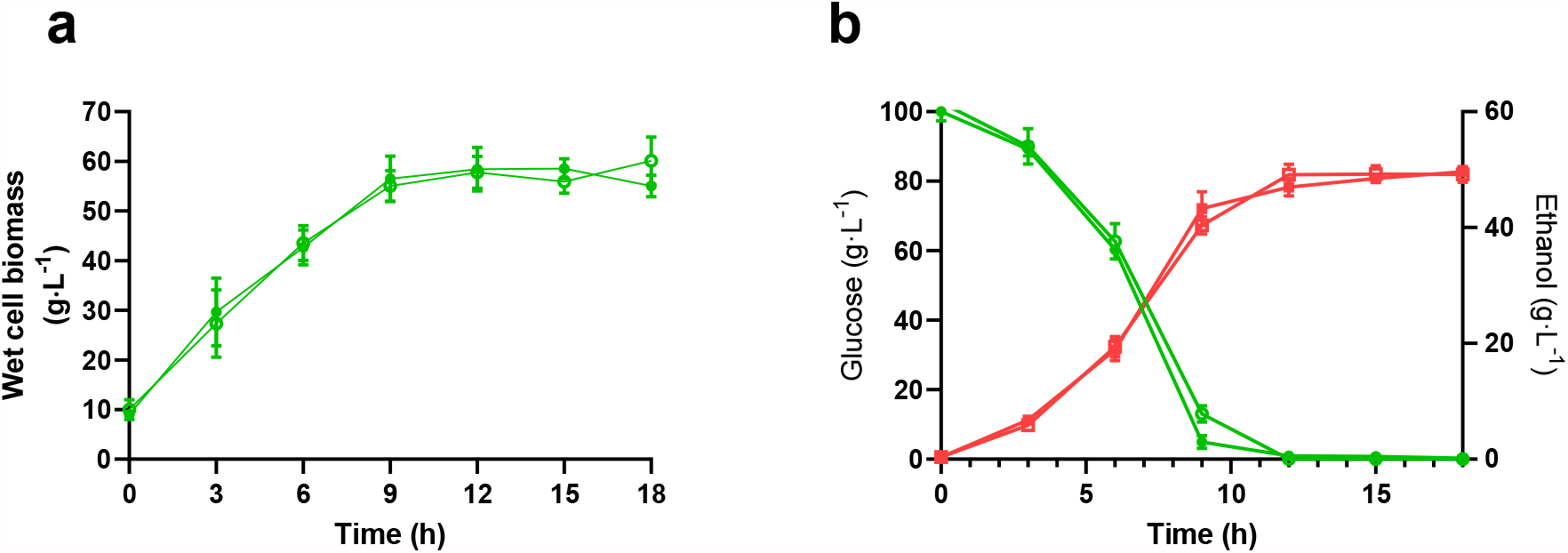
Kinetics of growth for the *S. cerevisiae* PE-2 strain and its isogenic *atg32* mutant in 0.5× YPD and 100 g·L^-1^ initial glucose under oxygen restricted conditions. **a**) wet biomass formation and **b**) sugar consumption and ethanol production. Datapoints represent the mean and the error bars the standard deviation of three biological replicates (n = 3). Filled circles represent the WT PE-2 and open circles represent *atg32 S. cerevisiae*.

### Fermentation performances of *S. cerevisiae* PE-2 and its corresponding *atg*32*Δ* mutant in a mimicked scaled-down biorefinery do not differ

Shiroma and co-workers (16) reported that blocking mitophagy improved the cell viability and ethanol titres during simulated Japanese Ginjo sake fermentation. To investigate whether the same strategy could improve fermentation performance under bioethanol production conditions encountered in Brazilian sugarcane biorefineries, triplicate cultures of *S. cerevisiae* PE-2 and of its isogenic *atg32Δ* mutant were performed using diluted molasses (obtained directly from a sugarcane refinery) as the cultivation medium, containing 140 g·L^-1^ of total sugars, in a scaled-down system that mimics the industrial process (cultivation method 2)(5). Both the WT and the *atg32Δ* strains behaved identically with respect to CO_2_ production - with a maximum CO_2_ of 2.0 ± 0.0 g (WT) and 2.0 ± 0.1 (*atg32Δ*) (Fig. 4a), reaching a relative ethanol yield of 92.5 ± 4.4 % (WT) and 94.0 ± 3.5 % (*atg32Δ*) of the maximum stoichiometric value of 0.511 g ethanol/g hexose, respectively (Fig. 4b), during five rounds of cell recycling including acid wash between two consecutive rounds. The final ethanol concentration reached was similar (WT: 56.0 ± 2.0 g·L^-1^; *atg32Δ*: 56.7 ± 1.6 g·L^-1^) and the cell viability remained extremely high through the five cycles for both strains (WT: 99.4 ± 0.3%; *atg32Δ*: 99.6 ± 0.4 %). The glycerol concentration remained similar for all the cycles (Fig. S1a, Supplementary Material) while yeast wet cell biomass was slightly higher in the WT strain along the cycles as compared to the *atg32Δ* mutant (Fig. S1b, Supplementary Material). However, the biomass increase, or decrease was not significantly different when the WT and the *atg32Δ* mutant were compared. Cells accumulate trehalose under nutrient starvation (28). Differences in trehalose accumulation were negligible (Fig. S1c, Supplementary Material). The cell numbers at the end of each cycle were slightly higher for *atg*32*Δ* (565 ± 20 × 10^6^ cells·mL^-1^) throughout the five cycles tested compared to the WT (583 ± 20 × 10^6^ cells·mL^-1^) (Fig. S1d, Supplementary Material).

**Figure 4:**
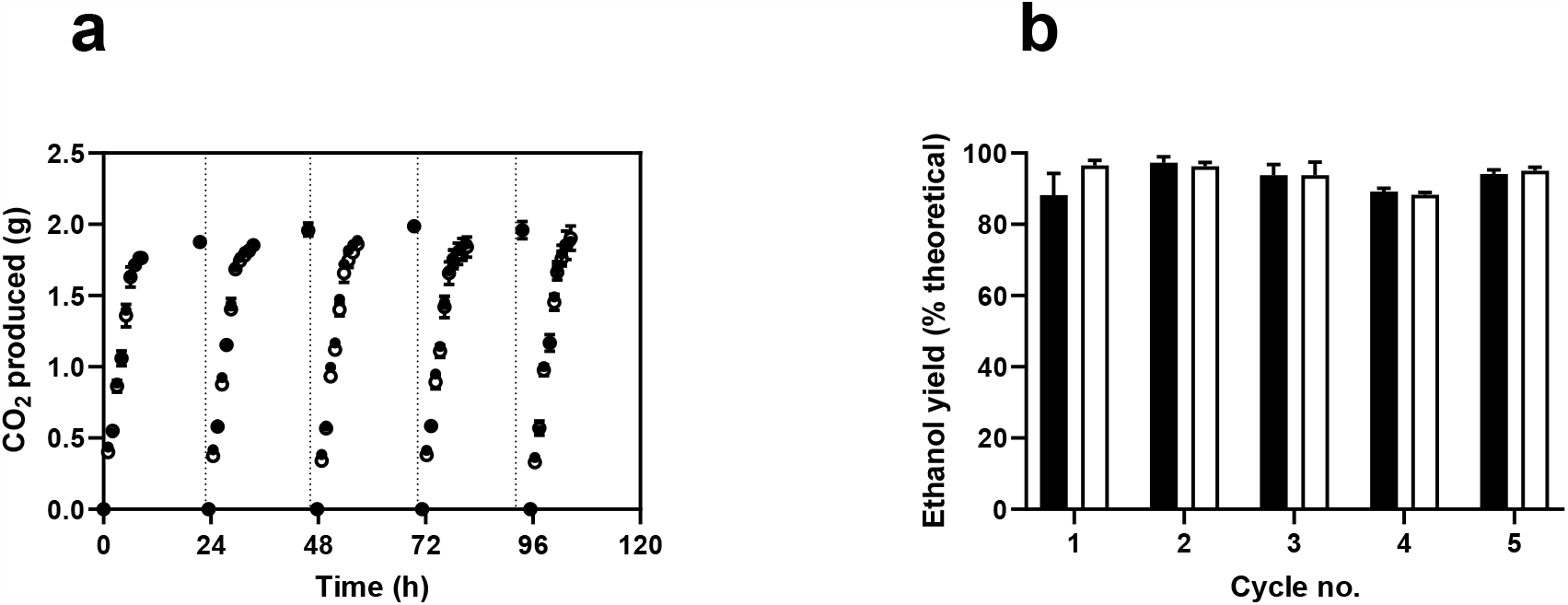
**a**) CO_2_ production and **b**) ethanol yields as % maximum for *S. cerevisiae* PE-2 (WT) and its corresponding *atg32* mutant in diluted molasses medium in a scaled down and mimicked process resembling Brazilian sugarcane fermentations. Filled circles/bars represent the WT PE-2 and open circles represent *atg32 S. cerevisiae*. Data represent the mean and standard deviation of three technical replicates and the experiment was performed twice.

Even though the absolute mass of wet cells (Fig. S1b, Supplementary Material) remained lower for PE-2 *atg32Δ* throughout the cell recycling process, the total cell concentration of the PE-2 *atg32Δ* strain was similar to that of WT. This is an indication of a reduced cellular volume for PE-2 *atg32Δ* as compared to the WT, which was corroborated by flow cytometry data that revealed a lower mean forward scatter (FSC) for the *atg32Δ* strain (Fig. S1e Supplementary Material). This observation agrees with the deletion of the *ATG32* gene in a sake strain background during the production of Ginjo sake (16). Nevertheless, the rest of the physiological parameters (CO_2_, glycerol, ethanol, biomass, and trehalose), including the ethanol yield, remained the same. It is likely that due to the low initial sugar concentration employed, when compared to what is used industrially, the final ethanol titre (57 g·L^-1^) achieved is lower than what is typically encountered in Brazilian mills. Because the cell viability remained extremely high at 98% throughout the five cycles, blocking mitophagy does not improve fermentation performance under conditions encountered during industrial fermentations for fuel ethanol production from sugarcane.

### Strain background does not influence the fermentation performance of *S. cerevisiae atg32* in a defined medium

As the deletion of the *ATG32* gene did not result in increased process performance in the *S. cerevisiae* PE-2 strain, the influence of strain background was investigated, to see whether this observation would hold for strain backgrounds different from PE-2. Because the cell recycling experiments are laborious, we decided to conduct the experiments in a synthetic medium (cultivation method 3) as described by Shiroma and co-workers (16). Cell viability and CO_2_ production were compared between the laboratory BY strain, industrial PE-2, and Ethanol Red *S. cerevisiae* together with their *atg32Δ* counterparts in SC medium using 150 g·L^-1^ glucose under static conditions. The CO_2_ profiles of the industrial WT strains of Ethanol Red, PE-2, and their *atg32* counterparts, were indistinguishable (Fig. 5a), while the laboratory BY strains had a lower rate of fermentation (Fig. 5a). Between days 0-2, volumetric CO_2_ production increased rapidly for PE-2 (29 ± 2 g·L^-1^·day^-1^), Ethanol Red (31 ± 4 g·L^-1^·day^-1^), PE-2 *atg32Δ* (33 ± 2 g·L^-1^·day^-1^), and Ethanol Red *atg32Δ* (33 ± 1 g·L^-1^·day^-1^) and then decreased to *ca*. 3.5 g·L^-1^·day^-1^ between days 3-11. On the other hand, BY WT and *atg32Δ* strains exhibited a steady CO_2_ production until day four (18 ± 1 g·L^-1^·day^-1^), 1.8-fold lower than the industrial strains, after which it began to decrease between days 5-11 to 2 g·L^-1^·day^-1^. The culture pH at the end of 10 days was ∼ 2.8 for all the strains. Cell viability was measured at the end of 11 days and PE-2 cells failed to produce viable colonies in a rich YPD medium (Fig. S2, Supplementary Material). However, the BY strains exhibited more CFUs compared to Ethanol Red strains. To see if low pH or high sugar concentration encountered during the fermentation affected growth, cells were plated on YPD at pH 2.5 and on 20% glucose. The WT and *atg32Δ* strains in all the three strain backgrounds exhibited remarkably similar growth physiology (Fig. S3, Supplementary Material) during spot assays on YPD and YP20D. PE-2 is known for its tolerance to low pH, as it has been selected for decades to withstand acid wash over multiple fermentation cycles in industry (6, 29). Ethanol Red WT and *atg32Δ* exhibited mild growth at the lowest cell concentration, and BY strains did not grow. We can conclude that *ATG32* deficiency did not improve the fermentation performance in a defined medium in all the three strain backgrounds.

**Figure 5:**
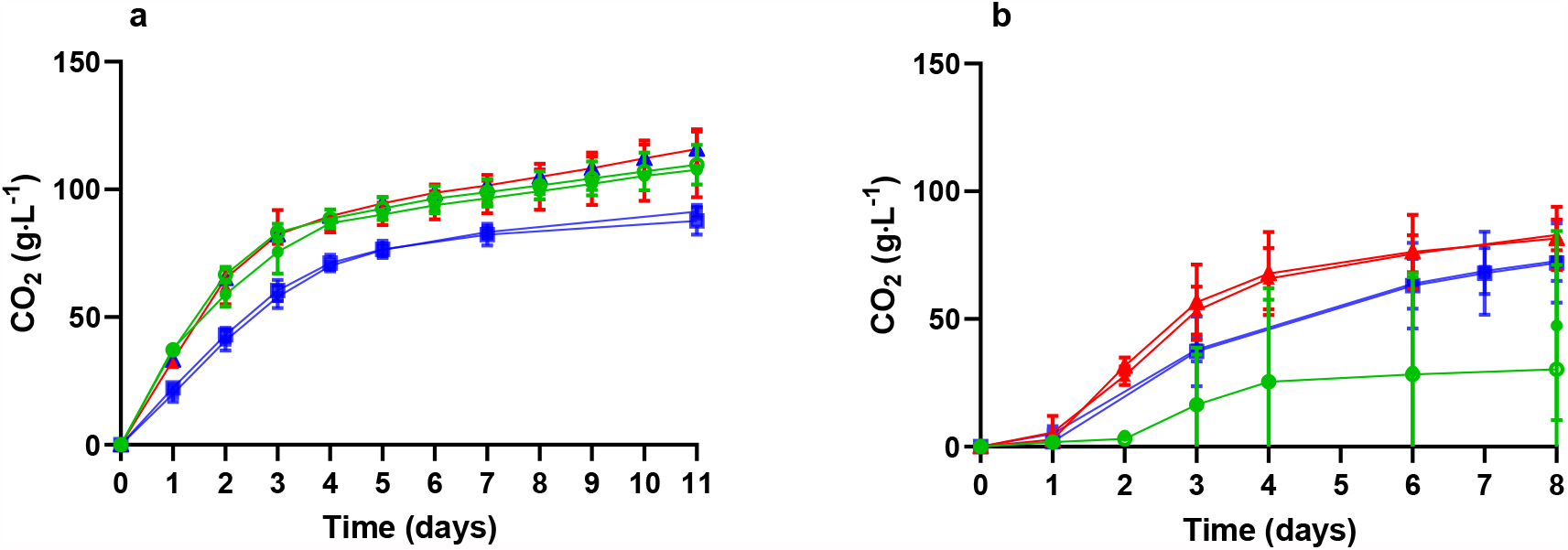
**a**) Time course of CO_2_ production for the WT and *atg32 S. cerevisiae* in SC medium using 150 g·L^-1^ glucose. **b**) Time course of CO_2_ production after re-pitching the pelleted cells from 11 days of fermentation with 40 mL of SC medium. Open markers represent the *atg32* strains and filled markers represent the WT strains; Ethanol Red is depicted in red, PE-2 is green, and BY in blue. Data represent the mean and standard deviation (n = 3).

### Ability to ferment after re-pitching is strain dependent

To test if the CFU counts reflected the metabolic vitality of cells in a liquid medium, cells from the end of fermentation were re-pitched with fresh SC medium for a second round of fermentation (30). Both Ethanol Red and BY strains were able to commence their fermentation, albeit at a lower rate than the first cycle (Fig. 5b) but only after an initial lag of one day. The Brazilian yeast PE-2 struggled to ferment, and there was a big variation in the CO_2_ production between the replicates, which is an indication of population heterogeneity, as demonstrated for acetic acid tolerance by Swinnen and co-workers (31). CO_2_ production was highest by day four after which it reached a plateau. Ethanol Red had the highest final CO_2_ production (WT: 83 ± 6 g·L^-1^; *atg32* : 82 ± 12 g·L^-1^), followed by BY (WT: 73 ± 8 g·L^-1^; *atg32* : 72 ± 16 g·L^-1^), and PE-2 (WT: 47 ± 37 g·L^-1^; *atg32* : 30 ± 41 g·L^-1^) respectively.

### Vast differences in the aerobic and oxygen restricted growth kinetics between laboratory and industrial strains

To investigate the dramatic differences in viability between the three strains in their WT backgrounds, they were cultured under oxic and oxygen restricted conditions in SC medium to monitor the viability kinetics over 10 days (cultivation method 4). During oxygen restricted conditions, the cell count increased between day 0 and 1, but once ethanol started to accumulate between days 1-6 the viable cell counts started to decrease steadily (Fig. 6a). The log_10_ decrease in CFU·mL^-1^ was 1, 1, and 0.4, for PE-2, Ethanol Red, and BY, respectively. However, between days 6-10, BY performed remarkably well, exhibiting a log_10_ decrease of 0.6, while for PE-2 and Ethanol Red, it had dropped further to 2 and 1.1, respectively. Glucose consumption was very rapid for PE-2 (35 ± 3 g·L^-1^·day^-1^) and Ethanol Red (32 ± 1 g·L^-1^·day^-1^) compared to BY (25 ± 1 g·L^-1^·day^-1^) (Fig. 6b). By day 5, all the glucose (150 g·L^-1^) was exhausted for PE-2, while for Ethanol Red, and BY, the residual glucose concentrations were ∼ 6 and 36 g·L^-1^ respectively (Fig. 6b). The residual glucose for BY after 10 days was ∼ 11 g·L^-1^. The final ethanol concentration reached were 79 ± 6, 79 ± 9, 72 ± 3 g·L^-1^ for PE-2, Ethanol Red, and BY, respectively (Fig. 6c).

**Figure 6:**
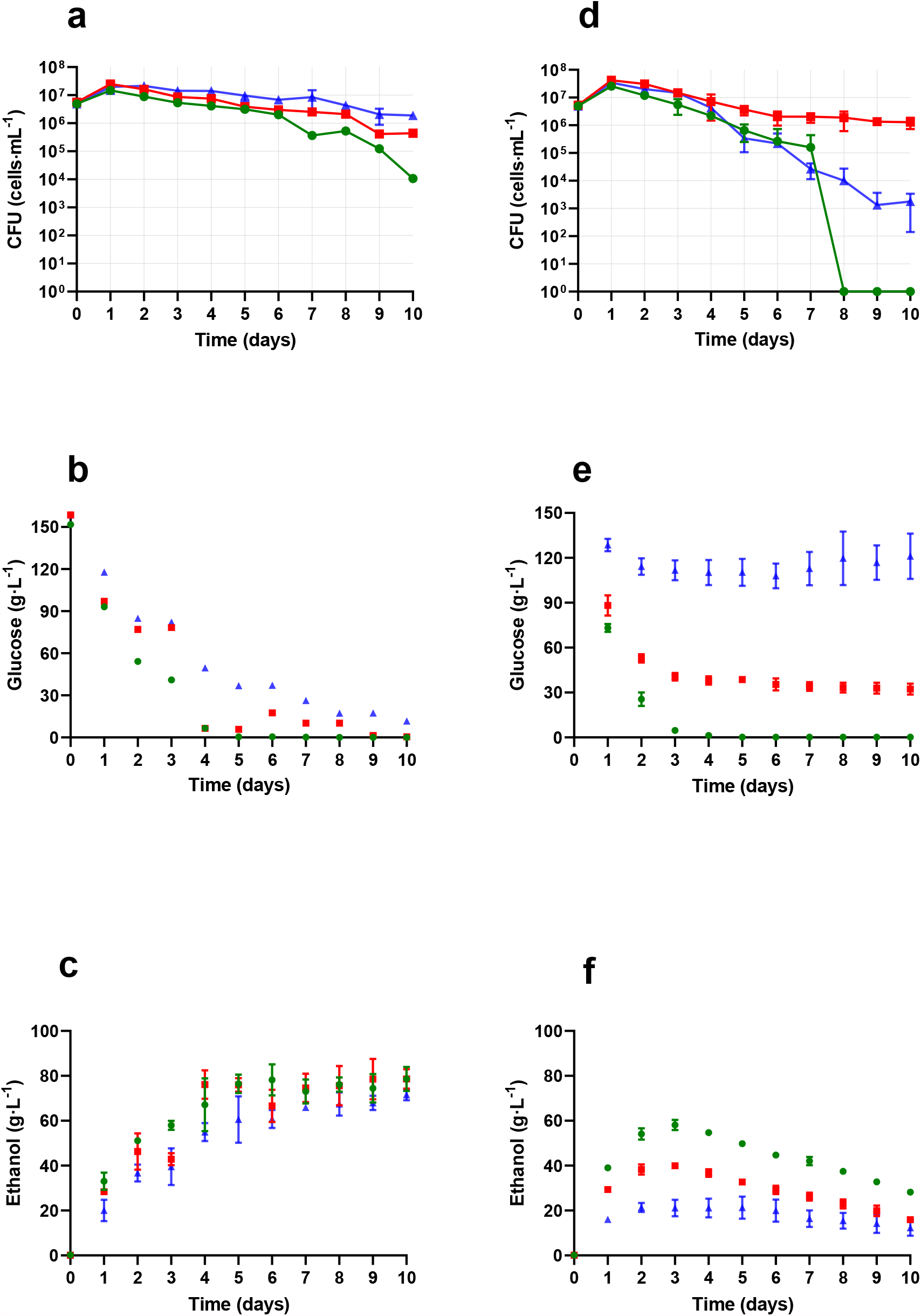
Growth kinetics of PE-2, Ethanol Red and BY yeast in SC medium under oxygen restricted and aerobic conditions. **Oxygen restricted**: **a)** CFU, **b)** glucose **c)** ethanol; **Aerobic**: **a)** CFU, **b)** glucose **c)** ethanol. Ethanol Red is depicted in red, PE-2 is green, and BY in blue. Data represent the mean and standard deviation (n = 3).

Unlike oxygen restricted conditions, the physiology of three yeast strains were remarkably different than under aerobic conditions. The viable cell counts decreased steadily between day 2-6 for PE-2 and BY and between days 8-10, no viable colonies were observed. Ethanol Red excelled among them and exhibited a log_10_ decrease in CFU of 1.5 between days 1-10 (Fig. 6d). PE-2 cells consumed glucose rapidly (68 ± 5 g·L^-1^·day^-1^) exhausting it by day 4 (Fig. 6e), while the rates were lower for Ethanol Red (51 ± 1 g·L^-1^·day^-1^) and BY (21 ± 2 g·L^-1^·day^-1^) (Fig. 6e). Also, glucose consumption stopped for Ethanol Red and BY from day 3, reaching a steady state glucose level of 34 ± 3 g·L^-1^ for Ethanol Red, and 115 ± 12 g·L^-1^ for BY (Fig. 6e). The final ethanol concentrations achieved were 58 ± 2, 40 ± 1, 23 ± 4 g·L^-1^ for PE-2, Ethanol Red and BY, respectively (Fig. 6f), lower than under oxygen restricted conditions owing to increased biomass formation due to respiro-fermentative metabolism.

## Discussion

High cell viability is vital for consistent cellular performance during bioproduction. The role of cellular housekeeping processes as a target for strain improvement during fermentation is a hitherto barely explored territory (15, 32–34). Shiroma and co-workers were the first in this regard, blocking mitophagy and observing improved ethanol formation (12% v/v for *atg32Δ vs* 11.75% v/v for WT BY4743) during Japanese sake fermentation (16). However, it should be pointed out that this improvement was only observed under conditions of Ginjo sake production and not under conditions of (conventional) sake production. Ginjo sake is produced under more severe nutrient limitation, when compared to conventional sake conditions, which is an indication that this limitation played an important role in the results obtained by Shiroma et al (16). The mechanisms by which the improvements are brought about have not been explored, but this increased ethanol production in the absence of mitophagy could be related to a lack of recycled nutrients normally provided by mitophagy, which cells use for biosynthesis. A combination of the absence of these internal building blocks and the concomitant excess of the external carbon and energy source – the sugar in the cultivation medium – would channel more of this carbon towards the fermentative pathway (36). In a different work, Jing and co-workers (35) report that deletion of *ATG32* not only decreased tolerance to ethanol stress in haploid BY 4742, but also led to a lower ethanol titre (7.25 %v/v) compared to the WT (7.5 %v/v) in standard YPD medium, which is nutrient rich. For their ethanol measurement, they relied on densitometry, just like Shiroma and co-workers, but this study used an enzymatic assay (16, 35). The improvement (or the decrease) reported in these studies are quite marginal (0.2% v/v), and it beckons us to question whether the accuracy of the measurement is reliable to claim physiological significance. It could also be that if a higher number of experimental replicates had been performed by Shiroma et al (16), no statistically significant difference would have been observed. In addition, the different analytical techniques employed prevents direct comparison between studies of any claims regarding the absolute ethanol titres. Morita and co-workers (36) report that deletion of *ATG32* increased the titres of 2,3-butanediol in engineered haploid BY4742 yeast, when compared to the parental strain. However, these authors did not observe higher ethanol yields or specific production rates in the *atg32* mutant, with respect to the values displayed by the parental strain, although deletion of *ATG32* significantly increased the maximum specific growth rate. It should be noted that these experiments were performed using a commercial synthetic yeast medium with 20 g/L initial glucose, meaning that nutrient limitation is much less prone to occur, when compared to the use of the same medium base, but with 150 g/L initial glucose, as performed by Shiroma et al (16, 35,36).

When mitophagy was blocked in two different industrial strains, which were tested for their performance in SC medium or using molasses in a scaled-down mimicked system, no statistical difference in ethanol titres or viability was observed in our study. Only marginal differences not related to fermentation performance were observed, such as a decreased cell size in the mutant strain, that was also reported by Shiroma and co-workers (16). This highlights the fact that benefits accrued under one set of conditions for one strain may not necessarily be shared by another strain in a different cultivation environment. This has been exemplified by Rodrigues and co-workers (37), who show that the maximum specific growth rate on sucrose is higher than on glucose for one wild yeast isolate, whereas this is not the case for other strains.

The three strains used in this study have vastly different origins. BY strain is a derivative of S288C, whose progenitor EM93-IC was isolated from rotting fig in 1938 (38). It is routinely propagated in the laboratory in YPD and SC medium. PE-2 was isolated from the Brazilian sugarcane fermentation industry (6). Ethanol Red is used extensively in the starch-based ethanol industry (39). The viability data from the fermentation experiment on SC medium (Fig 6a, 6d) corroborated the view that cells are adapted to their ecological niches and might fare poorly in environments that are not optimal for their growth.

The BY strain harbours a mutation in the *HAP1* gene (40) that renders it sensitive to aerobic growth. This could explain the retarded growth behaviour and the high residual glucose levels observed under aerobic conditions (Fig 6e). *HAP1* encodes a heme-dependent transcription regulator that controls the expression of genes involved in fermentation and ergosterol biosynthesis (41). BY cells performed very well under oxygen restricted conditions, possibly because of the absence of the repressor function of Hap1p (41, 42). BY strains which are derived from S288C are reported to be a phenotypic extreme and its widespread use as a norm for *S. cerevisiae* is a concern (43).

Despite these remarkable differences among the three strains investigated here, once mitophagy was blocked via deletion of *ATG32* in each background, no difference in fermentation performance was observed between each reference strain and its corresponding *atg32* mutant. This clearly indicates that the results reported by Shiroma and co-workers (16) in the context of Ginjo sake fermentation cannot be extrapolated to other strain backgrounds and/or conditions. Since we explored three different strain backgrounds and different cultivation conditions, we caution that the benefits of blocking mitophagy for ethanol formation in the context of Ginjo sake fermentations, if corroborated by future studies, are peculiar to those conditions and not necessarily the rule.

## Author contributions

VR carried out the strain characterisation and fermentation experiments; KPE, GCGC carried out the cell recycling experiments; BAW, DHMP, FZ constructed the knockout strains; EHH performed the Western blotting and fluorescence imaging. VR, TOB, AKG, EHH designed, and planned the experiments. VR wrote the manuscript, and it was approved by all the co-authors.

## Funding

We thank BBSRC-KE funding (R/160337) for VR and Dr. Mikael Molin (Department of Biology and Biological Engineering, Division of Systems Biology, Chalmers University of Technology, Sweden) for providing feedback. TOB would like to acknowledge *Fundação de Amparo à Pesquisa do Estado de São Paulo* (FAPESP) for grant 2018/17172-2. AKG would like to acknowledge *Fundação de Amparo à Pesquisa do Estado de São Paulo* (FAPESP) for grant 2017/08464-7 and *Conselho Nacional de Desenvolvimento Científico e Tecnológico* (CNPq) for fellowships 307266/2019-2.

## Supplementary Material

### Supplementary Figures

**Supplementary Figure S1:**
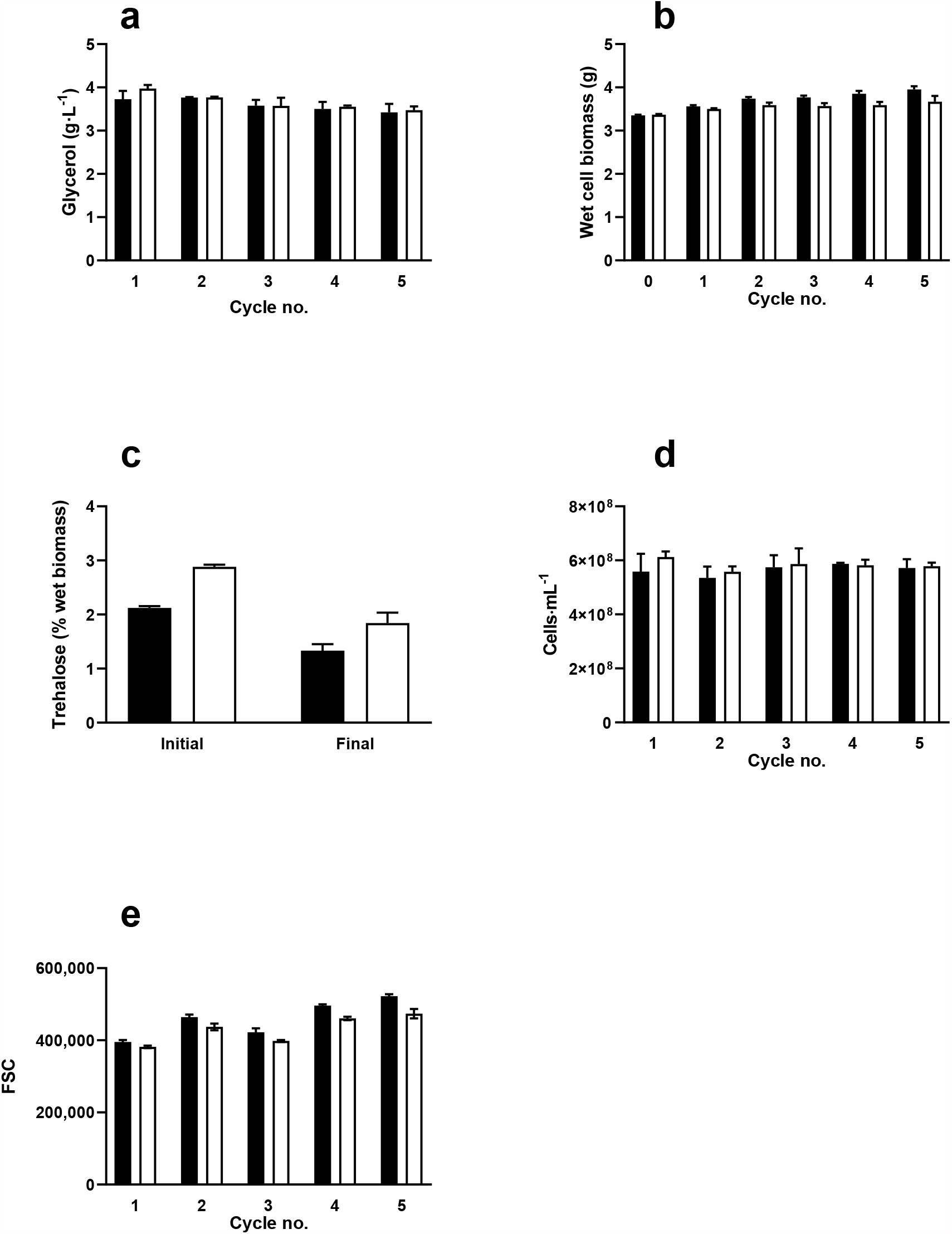
**a**) glycerol **b**) biomass **c**) trehalose **d**) cell number **e**) mean forward scatter (FSC) value for WT PE-2 and *atg32 S. cerevisiae* in diluted molasses medium in a scaled down and mimicked Brazilian fermentation process. Data represent the mean and standard deviation of three technical replicates, but the experiment was performed twice.

**Supplementary Figure S2:**
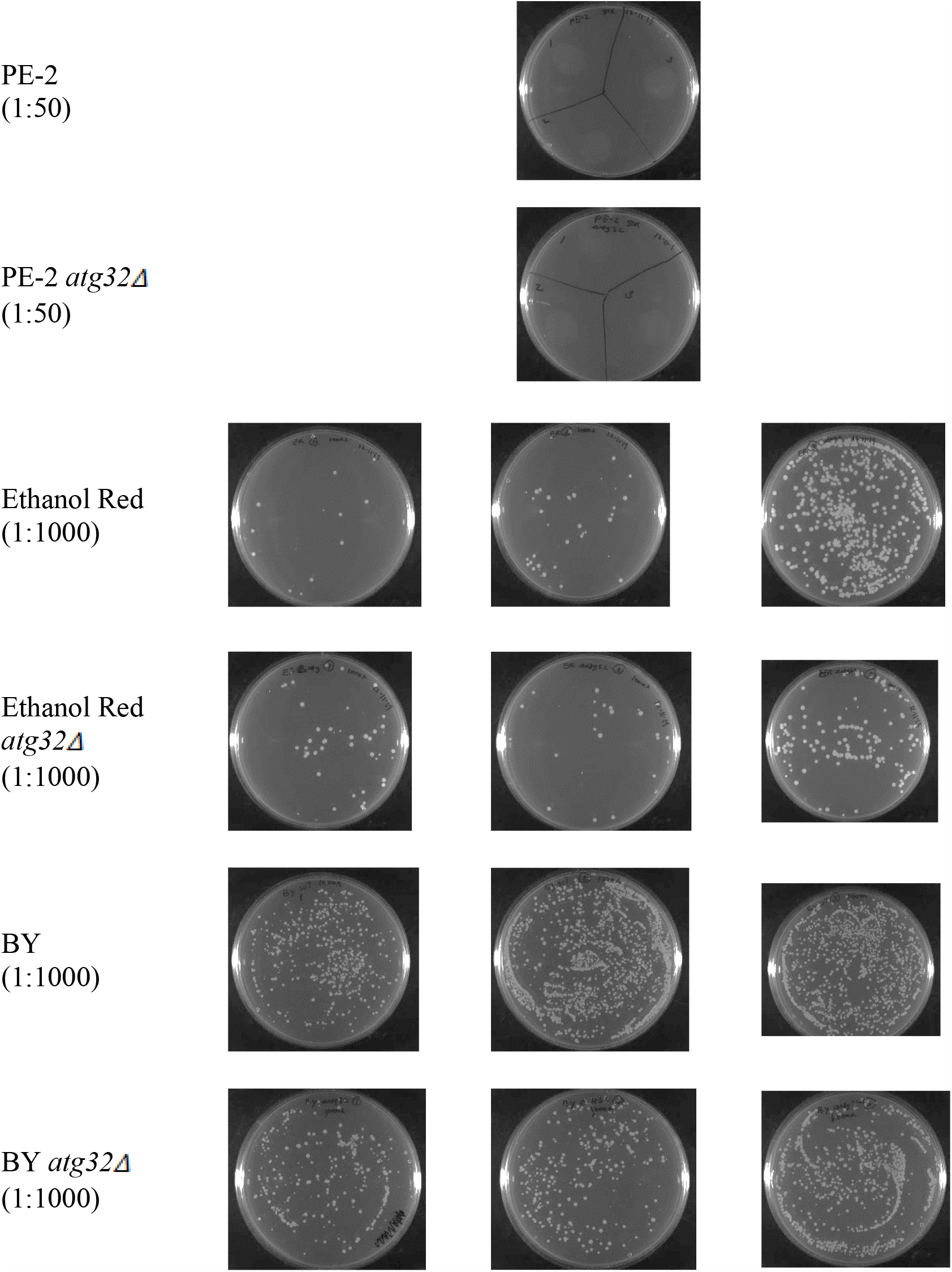
CFU of WT and *atg32 S. cerevisiae* in YPD after growth for 10 days in SC medium using 150 g·L^-1^ glucose for PE-2, Ethanol Red, and BY yeast.

**Supplementary Figure S3:**
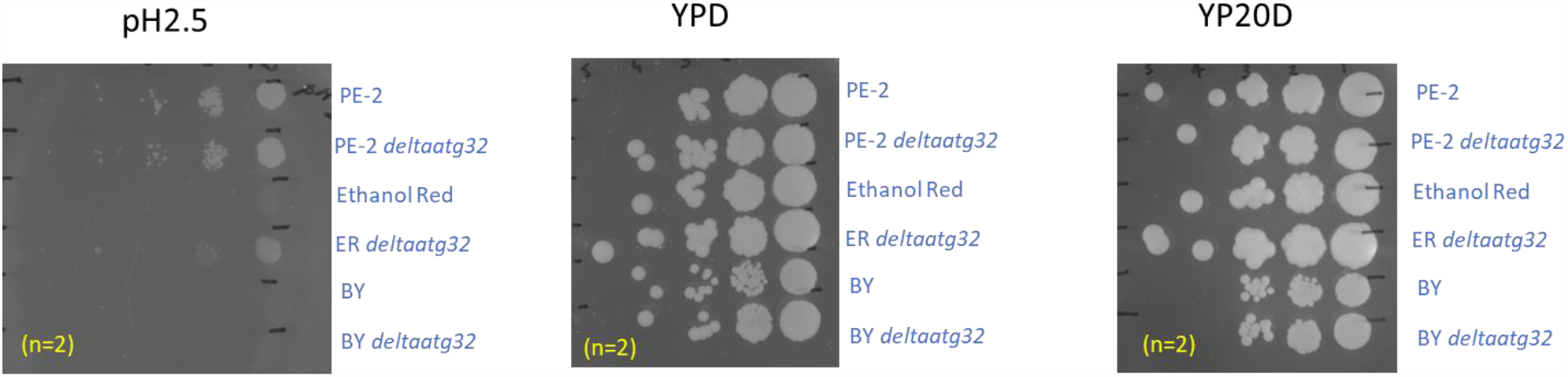
Spot assays of PE-2, Ethanol Red, and BY yeast in pH* 2.5, 2% YPD and 20% YPD. (n = 2). (*For making pH 2.5 YPD plates, the volume of sulphuric acid needed to bring the pH of a known volume of standard YPD to 2.5 was measured; the acid solution was filter sterilised and then aseptically added to YPD agar just before pouring onto plates.)

### Supplementary tables

**Table S1:**
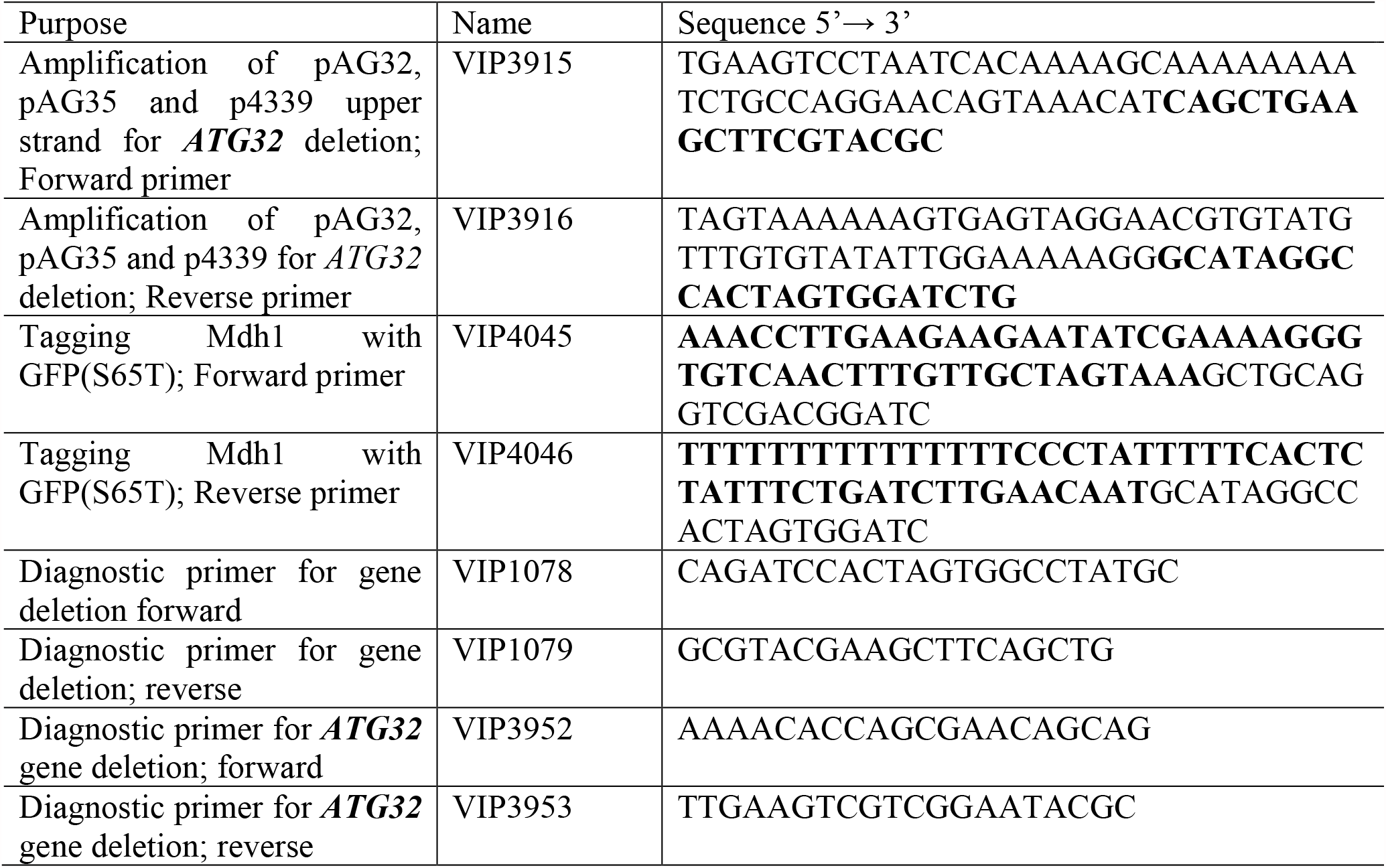
Primers used in this study.

**Table S2:** List of plasmids used in this study.

| Name | Relevant characteristics | Source |
| --- | --- | --- |
| pAG32 | HphMX | Euroscarf |
| pAG35 | NatMX | Euroscarf |
| p4339 | NatMX | (21) |
| pFA6a-GFP(S65T)-KanMX | GFP KanMX | (23) |
| pFA6a-GFP(S65T)-HIS3MX6 | GFP His3 | (23) |

